# Development of fluorescent and affinity tagged variants of rotavirus nonstructural protein 4 using the reverse genetics system

**DOI:** 10.64898/2026.09.04.749488

**Authors:** Ethan M. Huleatt, Abby L. Lipinski, Jacob L. Perry, Sue E. Crawford, Mary K. Estes, Joseph M. Hyser

## Abstract

Rotavirus (RV) nonstructural protein 4 (NSP4) is a multifunctional viroporin and enterotoxin that also serves an essential structural role in virion assembly, making it both a central virulence factor and a potential therapeutic target. However, studying NSP4 within infectious virus has been hindered by essential RNA packaging signals at the 5′ end of gene segment 10 (gs10) and an DLP-binding domain at its C-terminus, both of which constrain conventional mutagenesis. Using a plasmid-based reverse genetics system, we generated six recombinant rotaviruses (rRVs) encoding tagged or reporter-fused NSP4 via three strategies: expression from an alternative segment (gs7) in a bicistronic arrangement, N-terminal tagging of gs10 with translation shifted downstream of the native packaging sequence, and retention of the gs10 5′ UTR and fusion of the first 20 amino acids fused to a reporter followed by bicistronic full-length NSP4 expression. All six rRVs were replication competent, with gs10-based constructs closely matching wild-type replication kinetics while gs7-based constructs showed modest attenuation and, in one case, genetic instability upon passage. Engineered NSP4 proteins were robustly expressed, correctly glycosylated, and capable of oligomerization. Live-cell imaging showed reporter fluorescence reliably tracked NSP4 synthesis and preceded NSP4-dependent intercellular calcium signals. Affinity purification of tagged NSP4 recovered viroplasm-associated and structural viral proteins along with candidate host interactors, including ANP32A, ANP32E, PPM1G, and H2AC1. These rRVs constitute a validated toolkit for dissecting NSP4 function during bona fide infection and establish a generalizable strategy for engineering constrained rotavirus gene segments.

## INTRODUCTION

Rotavirus (RV) is a double-stranded segmented RNA genome virus within the Reoviridae family and is recognized as one of the most important etiological agents for acute gastroenteritis in infants and young children worldwide(1, 2). The segmented nature of its genome, which comprises eleven distinct RNA gene segments (gs), allows for both modular genetic organization and the potential for reassortment between different strains, a feature that has been exploited in the development of vaccines but also complicates molecular studies (3). RV infection is characterized clinically by profuse watery diarrhea, vomiting, and fever, and, in severe cases, can lead to dehydration and death (1). While the introduction of live attenuated RV vaccines has significantly reduced the global burden of disease, these vaccines show reduced efficacy in resource-limited settings, where disease incidence remains high (4, 5). This discrepancy underscores the need for a deeper understanding of the molecular biology and pathogenesis of RV, both to inform improvements in current vaccination strategies and to identify alternative therapeutic targets.

These gaps are particularly evident in studies of RV nonstructural protein 4 (NSP4), one of the virulence factors for RV pathogenesis (6). NSP4 is a multifunctional glycoprotein which localizes to the endoplasmic reticulum (ER) encoded by gene segment 10 (gs10) (7, 8). On the one hand, NSP4 functions as both a viral enterotoxin and a viroporin (9–11). Its expression perturbs intracellular calcium homeostasis; a process linked to the activation of chloride secretion in intestinal epithelial cells and the subsequent onset of secretory diarrhea (9). NSP4 viroporin-induced disruption of calcium signaling also triggers intercellular calcium waves (ICWs), which propagate signaling changes to neighboring uninfected cells, amplifying the diarrheal response (12). This signaling induces a transcriptional signature characteristic of interferon-independent innate immune activation indicating NSP4 as playing a role in host pathogen recognition (13). On the other hand, NSP4 plays a structural role in the viral life cycle. NSP4 colocalizes and supports the development of mature viroplasms, along with the core viroplasm proteins NSP2 and NSP5 (14). It interacts directly with the structural proteins VP4, VP6 and VP7, facilitating the budding of intermediate double-layer particles (DLPs) to acquire the outer capsid of the triple-layer particles (TLPs), a step essential for producing infectious virions (15–17). These dual functions, as virulence factor and an essential component of progeny virus assembly, make NSP4 both a subject of fundamental virology research and a potential therapeutic target.

Although NSP4 is central to RV biology, significant experimental limitations have hampered mechanistic studies dissecting its structure and function. Chief among these is the difficulty of manipulating gs10 without impairing the virus’s ability to package its genome correctly or produce a functional NSP4 protein. The first approximately 100 nucleotides of gs10 are particularly problematic. This sequence spans both the 5′ untranslated region (UTR) and the beginning of the NSP4 open reading frame (ORF) (Fig. 1A), and evidence from both RV and other members of the *Reoviridae* family indicates that it contains critical RNA elements required for genome packaging (18, 19). Disruptions in this region can lead to profound defects in virus replication (20). In addition, because this sequence also encodes the N-terminal signal peptide that targets NSP4 to the ER and key glycosylation sites, modifications in this region risk impairing NSP4’s correct localization and subsequent folding.

**Figure 1.**
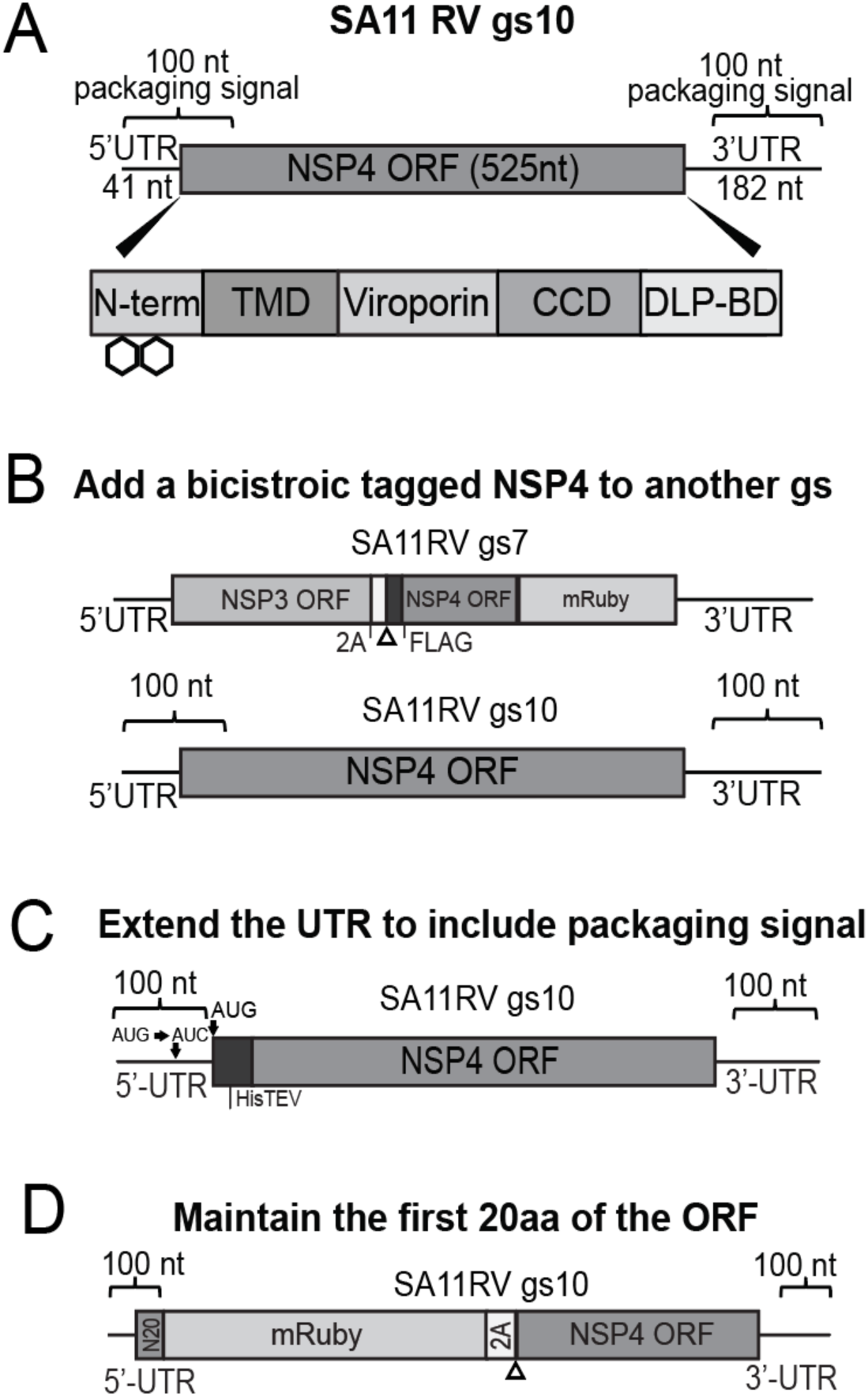
Overview of NSP4 tagging strategies. (A) Diagram of RV gs10 demonstrating both UTRs and the NSP4 ORF with the N-terminus domain, transmembrane domain (TMD), viroporin domain, coiled-coil domain (CCD) and DLP-BD. (B) Diagram of NSP4 tagging approach involving the expression of NSP4 with a marker from gs7 via a bicistronic arrangement. (C) Diagram of NSP4 tagging approach involving N-terminal tagging of NSP4 by shifting the ORF downstream of the native packaging sequence and mutating the original start codon. (D) Diagram of NSP4 tagging approach involving retention of the first 20 amino acids of the NSP4 ORF fused to a reporter and followed with a bicistronic arrangement to encode the complete NSP4 ORF.

The other major constraint lies at the opposite end of NSP4. The C-terminal domain of NSP4 contains the DLP-binding domain (DLP-BD) (Fig. 1A), which mediates the interaction between NSP4 and assembled DLPs (17, 21). This interaction is indispensable for the proper acquisition of the outer capsid layer of the TLP and for maintaining replication fitness. While some experimental truncations of the DLP-BD were tolerated in one virus strain (SA11), these truncations significantly reduced viral yields, indicating that even partial perturbation of this domain can compromise the efficiency of the viral life cycle (22).

These challenges have prevented systematic mutagenesis of NSP4 in the context of infectious virus. Past studies instead used overexpression systems to express NSP4 variants as recombinant NSP4 fragments (11–13, 23). While such approaches have been informative for elucidating certain biochemical and structural properties, they lack the physiological context of viral infection, where protein function is tightly integrated with the spatial and temporal dynamics of replication (22).

The recent development of plasmid-based reverse genetics systems for RV have enabled precise engineering of the RV genome, allowing for the targeted modification, replacement, or insertion of viral gene segments (24). This system allows the recovery of infectious recombinant RV (rRV) entirely from cDNA clones, enabling precise manipulation of individual genome segments. The technique involves the co-transfection of cells with plasmids encoding each of the eleven RV genome segments under the control of a T7 promoter, alongside support plasmids encoding helper proteins needed for transcription, capping, and replication (24). The result is a powerful platform for targeted genetic engineering of RV, permitting the insertion of epitope tags, reporter genes, or targeted mutations within specific genome segments.

The reverse genetics platform has facilitated the production of chimeric rRVs in which one or more genome segments were replaced with corresponding segments from a heterologous strain (13). Researchers have succeeded in generating bicistronic genome segments by introducing self-cleaving peptide sequences (e.g. P2A or T2A) between two coding regions, enabling the expression of two separate proteins from a single gs (25, 26). Bicistronic constructs have been used to tag NSP1 (gs5) and NSP3 (gs7) with fluorescent reporters, facilitating live-cell imaging of viral protein expression and dynamics without impairing virus replication (25, 27).

However, applying this strategy directly to gs10 has proven more difficult. The need to preserve both the 5′ packaging signal and the C-terminal DLP-BD means that any tagging strategy must be carefully engineered to avoid interfering with either of these critical regions. This dual constraint has significantly limited the success of our previous attempts to generate rRVs bearing modified NSP4.

To address these barriers, we devised an approach that takes advantage of the flexibility of the reverse genetics system while accommodating the structural and functional constraints of gs10. Rather than relying on a single tagging strategy, we designed three configurations, each engineered to circumvent the problems posed by direct modification of the gs10 ORF at its most sensitive regions (Fig. 1). In our first approach, we expressed NSP4 from an alternative genome segment, such as gs7, in a bicistronic arrangement that appends either a fluorescent reporter such as RFP or a genetically encoded calcium indicator (e.g. GCaMP8s) to the C terminus of NSP4 (Fig. 1B). By moving expression of the tagged NSP4 to gs7, this design leaves gs10 intact, ensuring the preservation of the packaging signals while allowing functional analysis of tagged NSP4 in the context of viral infection. For our second approach, we generated N-terminally tagged NSP4 variants by shifting the ORF downstream of the native packaging sequence and introducing a point mutation to prevent translation initiation at the original start codon (Fig. 1C). This design retains the essential 5′ RNA elements for packaging while permitting the addition of short N-terminal fusions without disrupting ER targeting. In the third configuration, we retained the gs10 5′ UTR and the first twenty amino acids of NSP4, which encompass the critical packaging elements, and fused them to an RFP reporter, and followed this with a bicistronic arrangement encoding the complete NSP4 ORF (Fig. 1D). This allows the expression of RFP from gs 10 to serve as a reporter of NSP4 synthesis throughout the infection cycle and N-terminal affinity tagging of the full length NSP4.

By adopting this multi-pronged design strategy, we preserve the most essential RNA elements of gs10 and ensure that at least one pool of NSP4 molecules retains an unmodified DLP-BD, safeguarding viral replication competence. The resulting rRVs offer a unique opportunity to interrogate the structural determinants of NSP4’s function, its role in viral assembly, and its capacity to modulate host cell calcium signaling in real time. Furthermore, these tools establish a flexible experimental platform for studying RV-host interactions, paving the way for future mechanistic investigations and the potential development of NSP4 targeted interventions.

## RESULTS

### Generation of Recombinant Rotaviruses Encoding NSP4 Constructs

In order to investigate strategies for genetically modifying NSP4 without disrupting essential gs10 packaging and NSP4 DLP assembly functions, we employed the plasmid-based reverse genetics system to generate rRV SA11 strains incorporating bicistronic or extended constructs in either gs7 or gs10 (Fig. 1). In gs7-based designs, NSP3 served as the primary ORF followed by a P2A self-cleaving peptide and three tagged forms of NSP4: (i) N-terminally 6xHis-tagged NSP4 (gs7-HN), (ii) NSP4 C-terminally tagged with mRuby (gs7-NR), or (iii) NSP4 C-terminally tagged with the calcium sensor GCaMP8s (gs7-NG) (Fig. 2A) (28). This strategy essentially keeps the native gs10 and expresses a second tagged version of NSP4 either for affinity purification or fluorescence microscopy, with the main caveat that these viruses now contain two copies of NSP4. For gs10, we engineered three constructs: (i) an extended UTR His-tagged NSP4 (gs10-HN), (ii) a chimeric construct expressing an N-terminal NSP4_1-20_-mRuby fusion as a reporter followed by a P2A element and full-length NSP4 (gs10-N20RN), and (iii) a similar construct incorporating a 6xHis affinity tag on the downstream NSP4 (gs10-N20RHN) (Fig. 2B). This strategy keeps a single copy of full-length NSP4 but enables affinity purification of NSP4 with an optional RFP-reporter for gs10 translation. Following co-transfection with the full complement of SA11 and the C3P3 accessory plasmids (see Materials and Methods for details), we successfully recovered infectious rRV for all six configurations.

**Figure 2.**
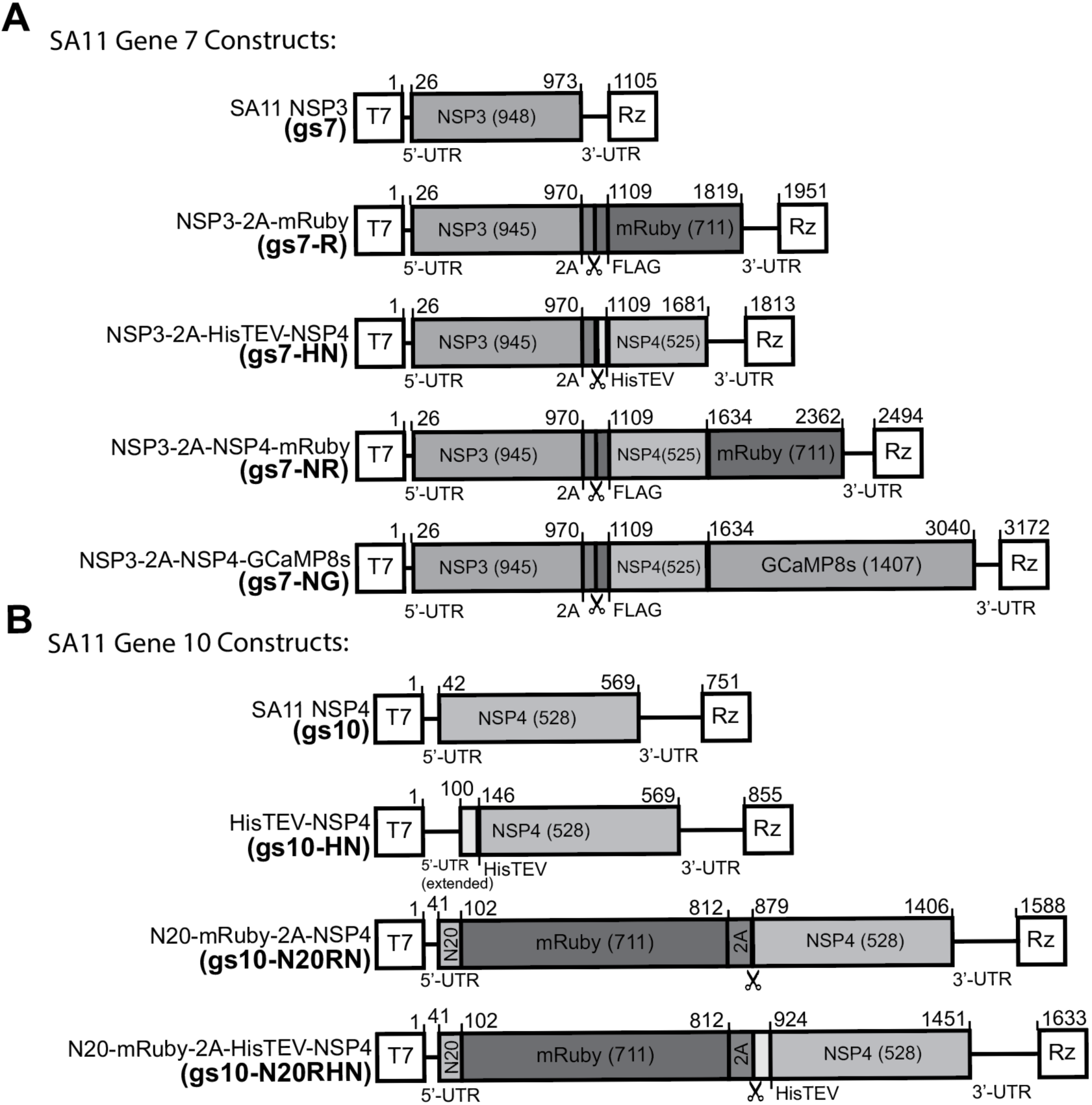
Schematic of recombinant rotavirus genome segment 7 and segment 10 NSP4 constructs. (A) Diagram of SA11 gs7 constructs used to generate rRVs. Constructs include the WT gs7 virus and derivatives encoding NSP3 linked by a 2A peptide to mRuby, His-TEV-tagged NSP4, FLAG-tagged NSP4-mRuby, or FLAG-tagged NSP4-GCaMP8s. (B) Diagram of SA11 gs10 constructs used to generate rRVs. Constructs include the WT gs10 virus and derivatives encoding an extended 5’ UTR His-TEV-tagged NSP4, N20-mRuby linked to NSP4 via a 2A peptide, or N20-mRuby together with His-TEV-tagged NSP4. Open reading frames, 5′ and 3′ UTRs, 2A cleavage sites, epitope tags, and reporter proteins are indicated. Numbers denote nucleotide positions within each genome segment. The designated short-hand name of each construct is bolded and shown in parentheses.

Electropherotype analysis shows results matching expected gs band sizes (Fig. 3A). Additionally, RT-PCR amplification, and sanger sequencing analysis confirmed incorporation of modified segments without evidence of gross rearrangements (Fig. 3B & Fig. S1). RT-PCR amplification and Sanger sequencing of gs7 and gs10 validated the integrity of engineered junctions. Of note, the gs10-HN rRV recombinant strain also contained the gs7-mRuby (gs7-R) segment that serves as a fluorescent reporter of infection/viral protein translation. Next, we performed serial passaging of the gs7 constructs to assess their stability. Passage of SA11 gs7-HN and gs7-NR exhibited stability of gs7. In contrast, serial passage of gs7-NG demonstrated overt genetic instability with high levels of recombination by passage three (Sup Fig. 1A). Sequencing of plaques showed that the rearrangements eliminated regions of the GCaMP8s tag, disrupting the calcium biosensor function. This instability was not unexpected as the gs7-NG size of 3172 bp is nearly 3-times that of the native gs7 and previous studies with SA11 gs7 constructs expressing large transgenes exhibited similar instability (25, 29). Thus, while the recombinant SA11 gs7-NG “calcium biosensor” strain requires routine plaque purification to maintain the full-length gs7-NG construct, it will nevertheless be a useful strain for direct calcium measurements at subcellular sites containing NSP4.

**Figure 3.**
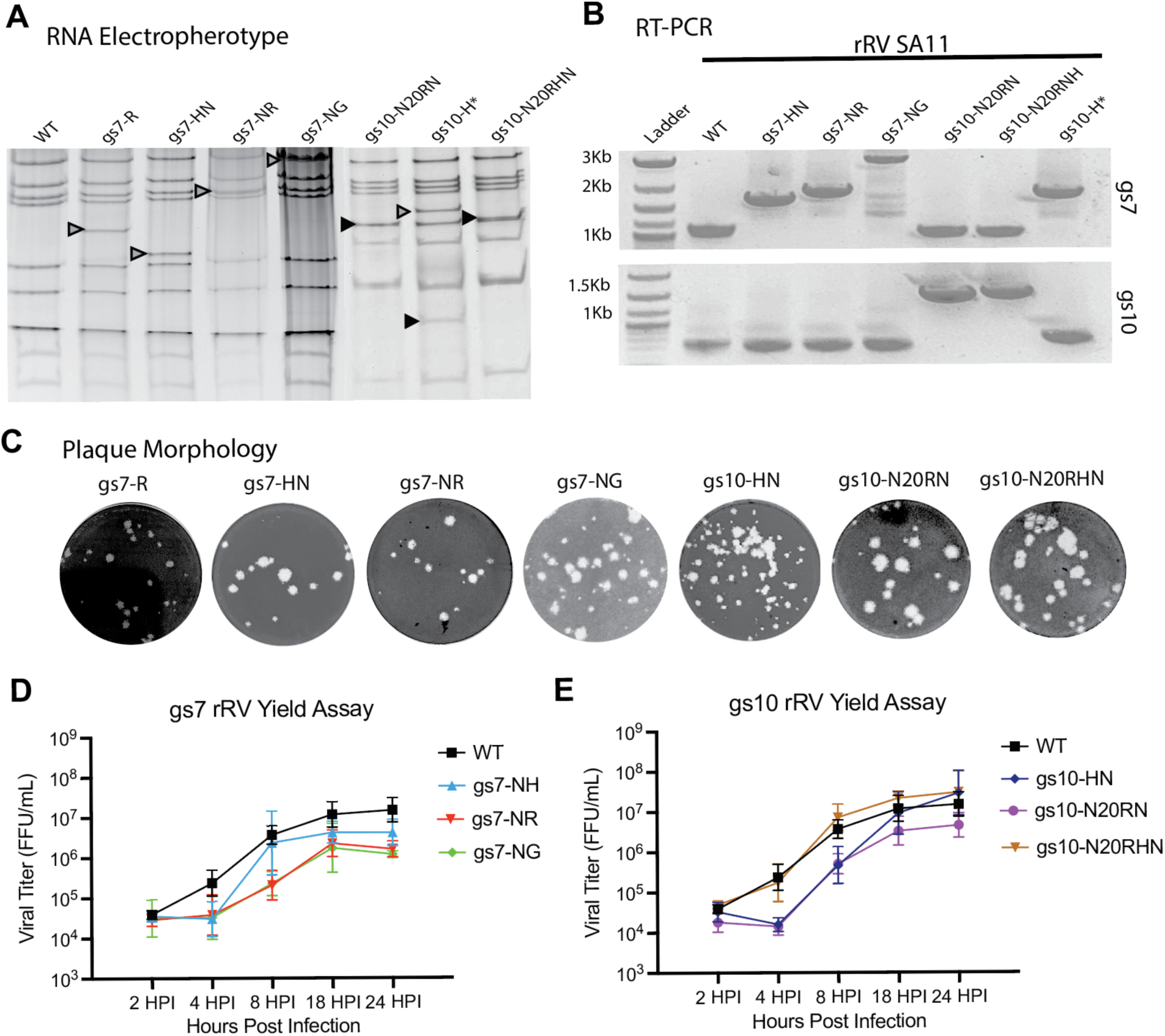
Generation and characterization of rRVs expressing tagged NSP4 variants. (A) Viral dsRNA genome electropherotypes of control rRVs (WT/gs7-R) and second passage rRVs encoding the indicated NSP4 variants. Arrowheads denote the modified genome segment containing the engineered NSP4 construct. (B) RT-PCR analysis confirming insertion of the engineered NSP4 sequences into gs10 rRVs or gs7 rRVs. Amplification of the modified genome segments produced products of the expected sizes and sequences. Primers utilized are listed in Table 1. (C) Representative plaque morphologies of recombinant viruses following infection of MA104 cell monolayers, demonstrating that all recombinant viruses formed plaques despite differences in plaque size and morphology. (D–E) Replication kinetics of rRVs. MA104 cells were infected and infectious virus production was quantified by focus-forming assay at the indicated hpi using anti-RV sera rabbit pool 42. (D) Replication kinetics of gs7 rRVs compared with the WT rRV. (E) Replication kinetics of gs10 rRVs compared with WT. Data are presented as geometric mean with geometric SD from 3 independent experiments.

**Table 1.**
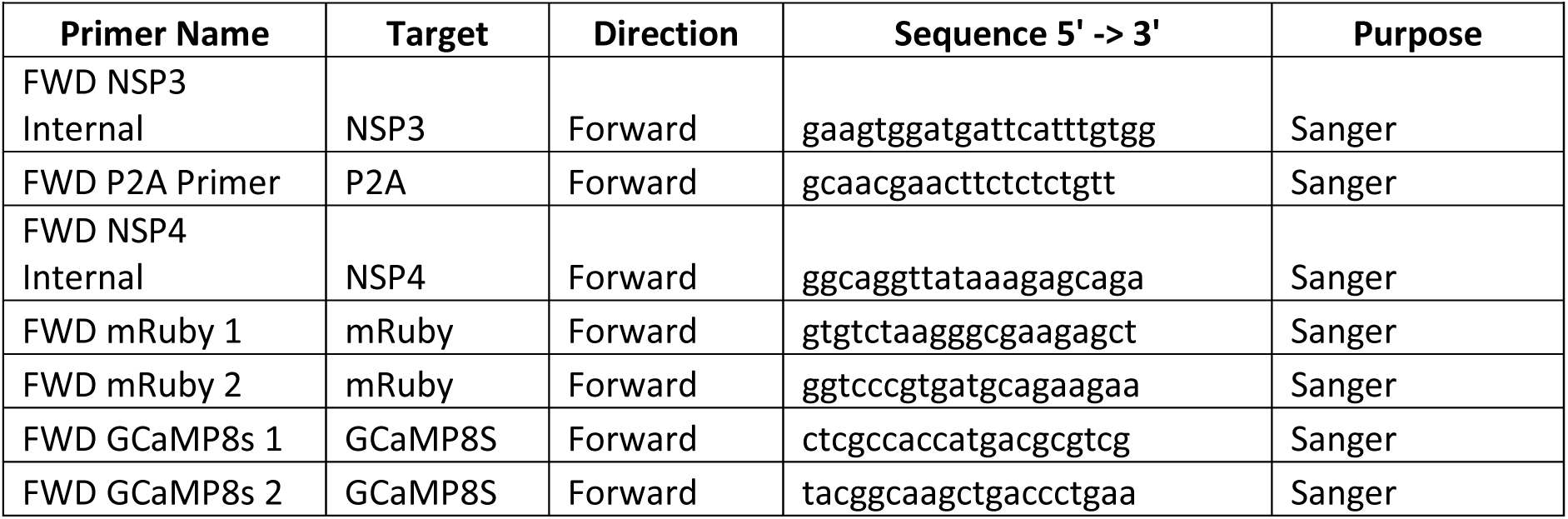
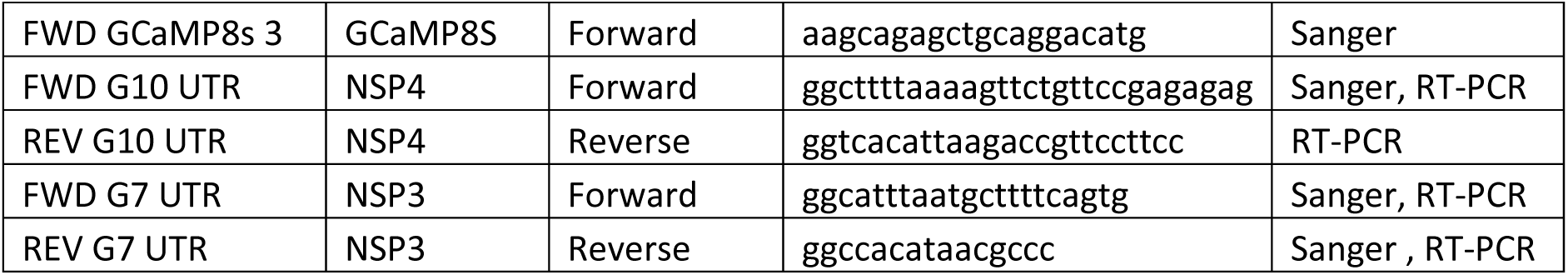
Primers utilized for RT-PCR and Sanger Sequencing.

### Replication Competence and Plaque Morphology

We evaluated our rRVs by plaque assay on MA104 cells to identify whether the engineering strategies of either gs7 or gs10 affected viral replication or spread. All rescued rRVs produced readily detectable plaques, demonstrating that the engineered genomes remained replication competent (Fig. 3C). The previously generated gs7-R reporter strain of SA11 was shown to have statistically insignificant differences in plaque morphology and yield compared to a recombinant wild-type (WT) SA11 (25, 30).

Among the rRV strains with engineered gs7, gs7-HN produced plaques that were generally larger and more uniform than the other gs7 constructs. gs7-NR exhibited an intermediate plaque phenotype, with plaques larger than gs7-R but smaller than gs7-HN. gs7-NG formed numerous plaques with variable diameters, including several relatively large plaques, suggesting that incorporation of the GCaMP8s reporter did not substantially impair cell-to-cell spread. However, we cannot rule out that the previously observed rearrangements in gs7-NG could have occurred during plaque formation, since it requires at least 3-4 rounds of replication and this could influence replication rate and spread.

Among the rRV strains with engineered gs10, all of the produced gs10 rRVs formed plaques comparable to gs7-R virus. The gs10-HN strain produced numerous small-to-medium plaques with morphology similar to gs7-R. Whereas gs10-N20RN (N20-mRuby-2A-NSP4) and gs10-N20RHN (N20-mRuby-2A-HisNSP4) produced plaques of intermediate size, with gs10-N20RHN appearing slightly more uniform than gs10-N20RN.

To evaluate whether modifications to gs7 or gs10 altered viral replication, we performed single-step growth curves in MA104 cells and quantified the infectious virus yield by focus-forming assays (Fig. 3D-E). Recombinant WT SA11 displayed the expected exponential increase in infectious titer between 4 and 18 hours post infection (hpi), reaching approximately 1-2x10^7^ FFU/mL by 24 hpi. All rRVs with an engineered gs7 exhibited slower replication kinetics than that of recombinant WT SA11 during the early phase of infection. At 8 hpi, titers of gs7-HN, gs7-NR, and gs7-NG remained below WT levels, indicating either delayed infection/entry stages of infection or slower initial production of infectious progeny. However, by 18 hpi, all gs7 rRVs demonstrated viral replication, reaching titers on the order of 10^6^-10^7^ FFU/mL. gs7-HN achieved the highest endpoint titer among the rRVs and most closely approached WT replication kinetics. In contrast, endpoint titers for both gs7-NR and gs7-NG were approximately one log lower than that of recombinant WT SA11. Endpoint titers for gs7-NR also exhibited greater variability between biological replicates, whereas replication of the gs7-HN strain exhibited more consistent yields. Finally, we similarly assessed the replication kinetics of the gs10 rRVs by single-step growth curves. In contrast to the gs7 rRVs, the gs10 rRVs displayed replication kinetics that more closely paralleled those of recombinant WT SA11. gs10-HN replicated nearly identically to WT throughout the time course and reached slightly higher endpoint titers by 24 hpi. Both gs10-N20RN and gs10-N20RHN showed a modest delay in replication during the early phase of infection, particularly at 4 hpi, but rapidly increased in titer between 8 and 18 hpi. By 24 hpi, all gs10 rRVs reached infectious titers that were within approximately 0.5-1-log_10_ of the recombinant WT SA11 control. Collectively, these data indicate that insertion of NSP4 fusion constructs into gs7 modestly attenuated viral replication, although all rRVs remained capable of producing high titer virus stocks. By contrast, insertion of mRuby and HisTEV-tagged NSP4 constructs into gs10 had minimal impact on viral growth, with rRVs displaying replication kinetics largely comparable to WT SA11.

### Expression of Tagged NSP4 Proteins during Infection

To determine the recombinant NSP4 expression kinetics, we infected MA104 cells with the indicated rRVs and collected lysates over a 10-hour time course. Viral protein expression was analyzed by immunoblotting using antibodies against NSP4, the FLAG epitope, the 6xHis tag, the 2A “self-cleaving” peptide, and a pooled anti-SA11 sera with gs7-R used as a positive control for detection of non-modified NSP4.

Initial characterization of the gs7-HN rRV confirmed successful expression and processing of the HisTEV-tagged NSP4 protein (Fig. 4A). Immunoblot analysis with an NSP4-specific antibody detected both the endogenous WT NSP4 and a slower migrating HisNSP4 species in infected cells. Expression of the recombinant protein was further confirmed by detection with anti-His and anti-2A antibodies, demonstrating our rRVs retained the inserted HisTEV sequence during viral replication. As expected, antibodies against SA11 proteins detected accumulation of major viral proteins, including VP2 and VP6 during infection, confirming productive viral replication.

**Figure 4.**
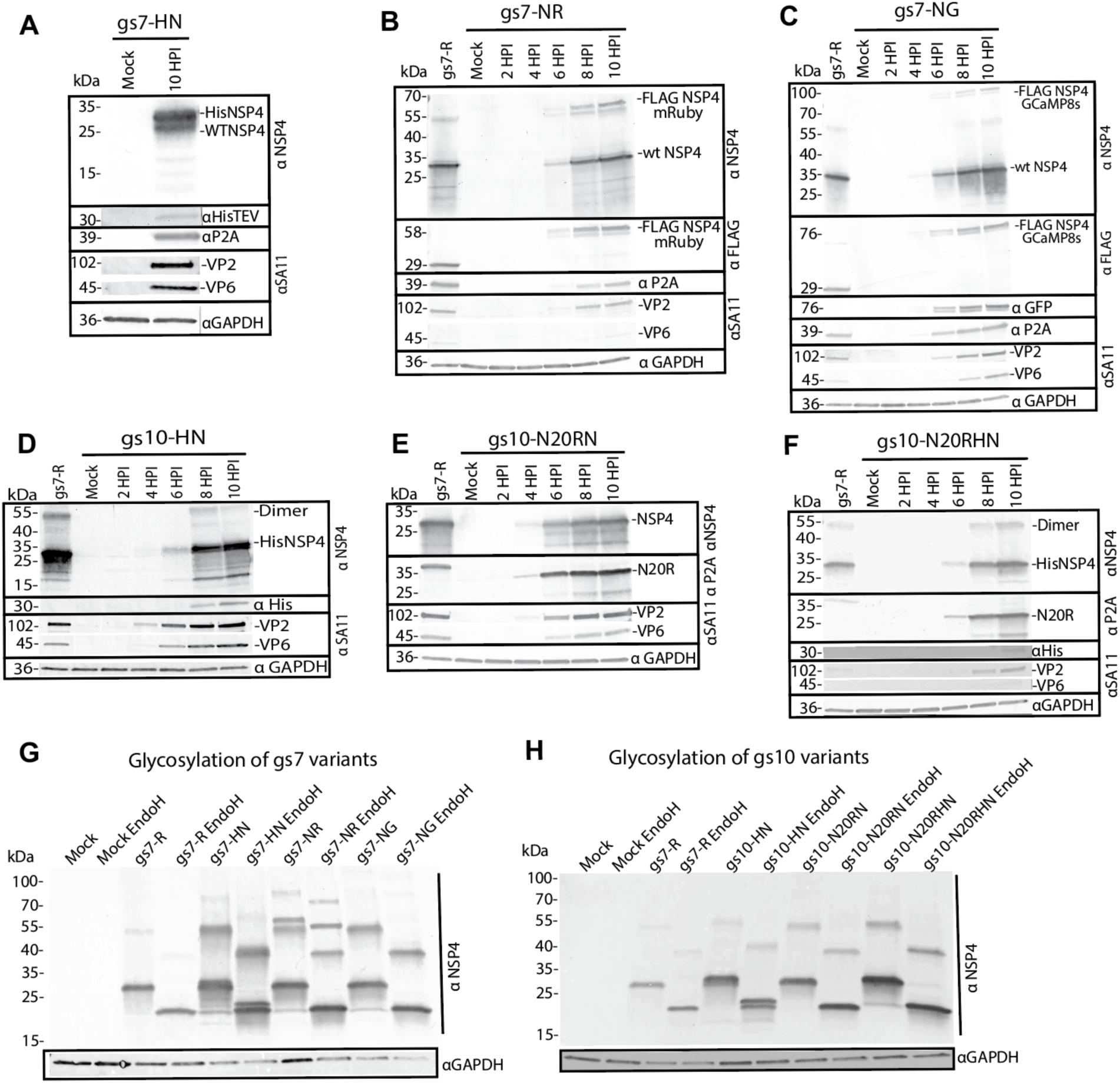
Expression and glycosylation analysis of recombinant rotaviruses encoding tagged NSP4 variants. (A–F) Time-course analysis of NSP4 expression during infection with recombinant SA11 rotaviruses expressing tagged NSP4 variants. MA104 cells were mock infected or infected with the indicated recombinant viruses and lysates were collected at the indicated hpi. Proteins were analyzed by immunoblot using antibodies against NSP4 (aa120-147) or the indicated epitope tag (His, FLAG, or P2A), together with sera against SA11 (PD30) to monitor viral protein expression. GAPDH served as a loading control. Each panel shows separate blots for each protein target analyzed, each with its own molecular weight marker. (A) gs7-HN, (B) gs7-NR, (C) gs7-NG, (D) gs10-HN, (E) gs10-N20RN, (F) gs10-N20RHN. The gs7-R was included as a control where indicated. (G–H) Glycosylation analysis of recombinant NSP4 variants by Endoglycosidase H (Endo H) digestion. Lysates from mock-infected or recombinant virus-infected cells were left untreated or treated with Endo H prior to immunoblotting with anti-NSP4. (G) gs7-derived NSP4 variants. (H) gs10-derived NSP4 variants. Blots are representative of 3 biological replicates.

We examined expression of the NSP4 fluorescent reporter fusions encoded from gs7 using the gs7-NR and gs7-NG rRVs (Fig. 4B-C). In cells infected with gs7-NR, anti-NSP4 immunoblotting detected both WT NSP4 (∼28 kDa) and the higher molecular weight FLAG-NSP4-mRuby fusion protein (∼55-60 kDa), which first become readily detectable between 4-6 hpi and increase through 10 hpi (Figure 4B). Anti-FLAG immunoblotting specifically recognized the mRuby fusion protein, while anti-2A antibody detected the residual 2A peptide sequence present on NSP3. Similarly, infection with gs7-NG resulted in progressive accumulation of the FLAG-NSP4-GCaMP8s fusion protein (∼90-100 kDa), which we detected by both anti-NSP4 and anti-FLAG antibodies beginning about 6 hpi and increasing through 10 hpi. In both FLAG rRVs, WT NSP4 continued to accumulate in parallel with the tagged proteins, indicating simultaneous expression from the native WT gs10. Expression of structural proteins (e.g., VP2 and VP6) paralleled accumulation of the recombinant NSP4 proteins, demonstrating that insertion of the reporter fusions did not interfere with the expression of other viral proteins (Fig 4B-C).

Our rRVs expressing tagged NSP4 from gs10 also exhibited robust protein expression throughout infection (Fig. 4D-F). Infection with gs10-HN produced a His-tagged NSP4 species that was readily detected by both anti-NSP4 and anti-His antibodies beginning at approximately 8 hpi (Fig. 4D). However, in the anti-NSP4 blot we detected an additional lower band. We believe that this smaller band results from an alternative translation initiation at the original start codon for NSP4, which this construct retained. Initiation from this start codon would produce an untagged, native NSP4 species. In addition to the monomeric protein, there is a double band higher molecular weight NSP4-reactive species consistent with an SDS-resistant NSP4 dimer, which is commonly seen for NSP4, and this indicates that the addition of a N-terminal HisTEV tag does not disrupt NSP4 oligomerization. Thus, immunoblot analyses indicate that this construct produces both the intended 6xHisNSP4 species and a native, untagged NSP4 species from the same gs10 construct in approximately a 1:1 ratio of tagged and untagged species.

The N-terminal mRuby fusion constructs encoded by gs10-N20RN and gs10-N20RHN likewise demonstrated efficient expression during infection (Fig. 4E-F). For gs10-N20RN, anti-NSP4 immunoblotting detected a single band of WT-sized NSP4, while anti-2A antibody identified the N20-mRuby-2A product that accumulated from 4 hpi onward. Similarly, the gs10-N20RHN virus expressed both the N20-mRuby reporter and the full-length His-tagged NSP4, as confirmed by anti-NSP4, anti-His, and anti-2A immunoblotting. Further, in this case there is only the single HisNSP4 species without any untagged native NSP4, which is the optimal set-up for affinity purification of NSP4 from a bona fide infection. Finally, consistent with the gs10-HN virus, the commonly observed, SDS-resistant NSP4 dimer accumulated during the late stages of infection.

We treated lysates from infected cells with endoglycosidase H (Endo H) to assess N-linked glycosylation to determine whether recombinant NSP4 proteins retained normal post-translational processing, (Fig. 4G-H) (8, 31). All recombinant NSP4 proteins exhibited a characteristic mobility shift following Endo H digestion, indicating that the engineered proteins remained glycosylated within the endoplasmic reticulum. Interestingly both NSP4 variants expressed from the gs10-HN rRV strain were Endo H sensitive, indicating both the native and 6xHis-tagged forms are ER targeted properly. Both gs7 and gs10 variants displayed Endo H-sensitive glycosylation comparable to WT NSP4, demonstrating that addition of fluorescent reporters, affinity tags, and peptide linkers did not disrupt glycosylation or ER targeting of NSP4.

Across all rRVs, expression of the engineered NSP4 proteins increased with infection kinetics and coincided with the accumulation of viral structural proteins, such as VP2 and VP6 (Fig 4). GAPDH levels remained constant throughout the infection, confirming equivalent sample loading. Collectively, these results demonstrate that recombinant NSP4 proteins containing fluorescent reporters, FLAG epitopes, or HisTEV affinity tags are robustly expressed during rRV infection while maintaining expression of the native gs10 NSP4 where appropriate. Furthermore, detection of higher molecular weight NSP4 species in multiple rRVs suggests that these engineered proteins retain the ability to form oligomeric complexes during infection, supporting their use for subsequent localization, biochemical, and protein interaction studies.

### Visualization of Reporter Expression and Calcium Dynamics

To determine whether reporter rRVs faithfully expressed fluorescent fusion proteins while retaining NSP4-mediated calcium signaling, we infected MA104-GCaMP6s cells with the indicated rRVs and analyzed NSP4 expression by fluorescence microscopy, immunoblotting, Endo-H digestion, and live-cell calcium imaging.

Representative fluorescence images demonstrated robust expression of all reporter constructs during infection (Fig. 5A-C). Cells infected with gs7-NR exhibited the expected strong ER associated mRuby fluorescence (Fig. 5A). Similarly, gs7-NG produced robust ER localized GCaMP8s fluorescence in infected cells (Fig. 5B), while gs10-NH, gs10-N20RN, and gs10-N20RHN displayed diffuse cytoplasmic mRuby fluorescence corresponding to the independently translated mRuby reporter (Fig. 5C). Together, these images confirmed efficient expression of each fluorescent reporter from the engineered genome segments and demonstrated that insertion of reporter proteins did not prevent productive infection.

**Figure 5.**
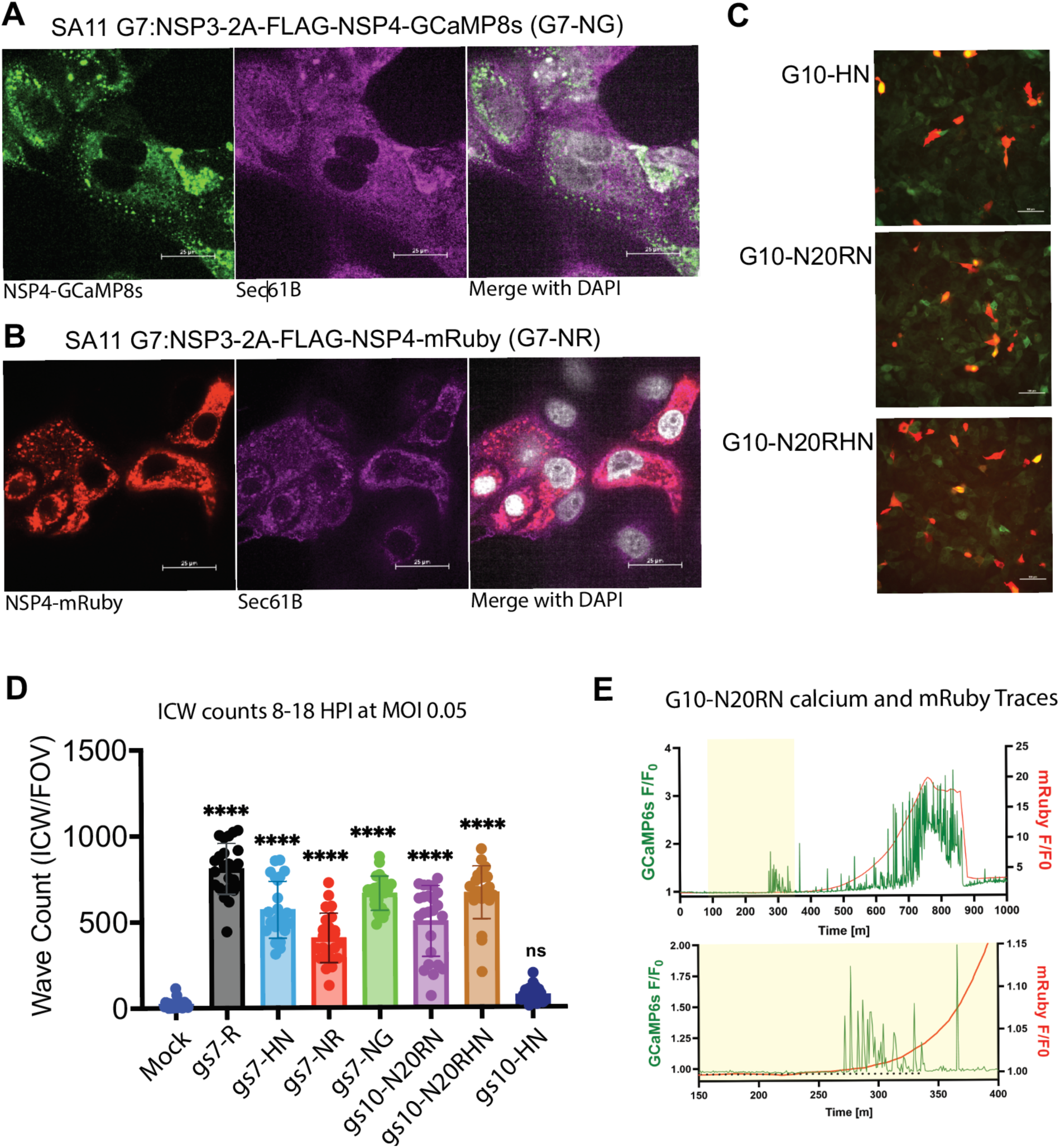
Visualization of reporter expression and calcium signaling in recombinant NSP4 reporter viruses. (A-B) Representative confocal microscopy images of MA104 cells infected with recombinant rotaviruses expressing gs-NG (green, calcium indicator) or gs7-NR (red), demonstrating tagged NSP4 expression during infection. Sec61 beta (ER marker) was stained for in far-red as well as DAPI (white). Scale bars = 25µM. (C) Representative images of MA104 cells infected with recombinant rotaviruses expressing gs7-R, gs10-N20RN, gs10-N20RHN demonstrating cytosolic reporter expression during infection. Scale bars = 100µM. (D) Quantification of ICW activity 8-18 hpi at an MOI of 0.05. ICW counts conducted by field of view (FOV) and compared among the indicated rRV and mock-infected cells. Data are presented as mean SD, with statistical significance determined relative to the mock; ****, P < 0.0001; ns, not significant. (E) Representative single-cell fluorescence intensity traces from cells infected with gs10-N20RN, showing GCaMP6s calcium dynamics (left y-axis) and mRuby reporter expression (right y-axis) over time. The expanded trace (bottom) illustrates the temporal relationship between increasing mRuby expression and the onset of NSP4-associated calcium signaling.

Because NSP4-induced calcium dysregulation is a hallmark of rotavirus infection, we next quantified ICWs generated by each rRV in MA104-GCaMP6s cells infected at an MOI of 0.05 (Fig. 5D). WT gs7-R virus produced robust calcium signaling characterized by approximately ∼800 ICWs per field of view (FOV). All reporter viruses expressing modified NSP4 from gs7 (gs7-HN, gs7-NR, and gs7-NG) generated significantly increased ICW counts compared to the mock infection. Likewise, the gs10 reporter viruses gs10-N20RN and gs10-N20RHN retained robust calcium signaling that was comparable to the WT. In contrast, gs10-HN failed to induce detectable calcium waves above mock-infected controls, indicating that insertion of the N-terminal HisTEV tag into the native NSP4 gene or the combinations of native and tagged NSP4 variants oligomerizing substantially impaired NSP4-mediated calcium signaling despite supporting productive viral replication.

We performed live-cell imaging using the gs10-N20RN virus in MA104-GCaMP6s cells to examine whether reporter fluorescence accurately reflected viral gene expression during infection (Fig. 5E). Simultaneous monitoring of GCaMP6s and mRuby fluorescence demonstrated that reporter expression progressively increased throughout infection and preceded the onset of robust calcium signaling. Expansion of the time frame representing the onset of increased calcium signaling revealed that the initial increase in mRuby fluorescence strongly coincided with the first transient calcium spikes. These observations indicate that insertion of a fluorescent protein reporter in gs10 generates a reliable real-time marker of gs10 protein expression, enabling tracking of the temporal relationship between NSP4 expression and calcium dysregulation.

Together, these results demonstrate that the reporter rRVs efficiently express fluorescent NSP4 fusion proteins or linked fluorescent reporters while maintaining normal NSP4 glycosylation. Furthermore, except for the gs10-HN construct, all engineered viruses preserved the characteristic NSP4-dependent calcium signaling phenotype, establishing these rRVs as valuable tools for simultaneously monitoring viral protein expression and calcium dynamics during rotavirus infection.

### NSP4–NSP4 Interactions Revealed by Pulldown and Mass Spectrometry

To establish whether tagged variants of NSP4 expressed from gs7 are incorporated into oligomeric complexes with native WT NSP4 expressed from gs10, rRVs expressing FLAG-tagged gs7 NSP4 constructs were subjected to anti-FLAG affinity purification followed by immunoblot analysis. As expected, we observed no enrichment of viral proteins in mock-infected controls (Fig. 6A). In contrast, FLAG pulldown of the gs7 rRVs resulted in recovery of multiple higher molecular weight NSP4-containing species in the elution fractions, demonstrating efficient capture of the tagged proteins. Importantly, we detected protein bands corresponding to both the tagged recombinant NSP4 and the lower molecular weight WT NSP4 species in the elution fractions of the gs7-NG and gs7-NR viruses, indicating that recombinant and WT NSP4 molecules co-purify within the same protein complexes. These findings demonstrate that NSP4 molecules expressed from separate genome segments retain the ability to assemble into mixed oligomeric complexes during infection.

**Figure 6.**
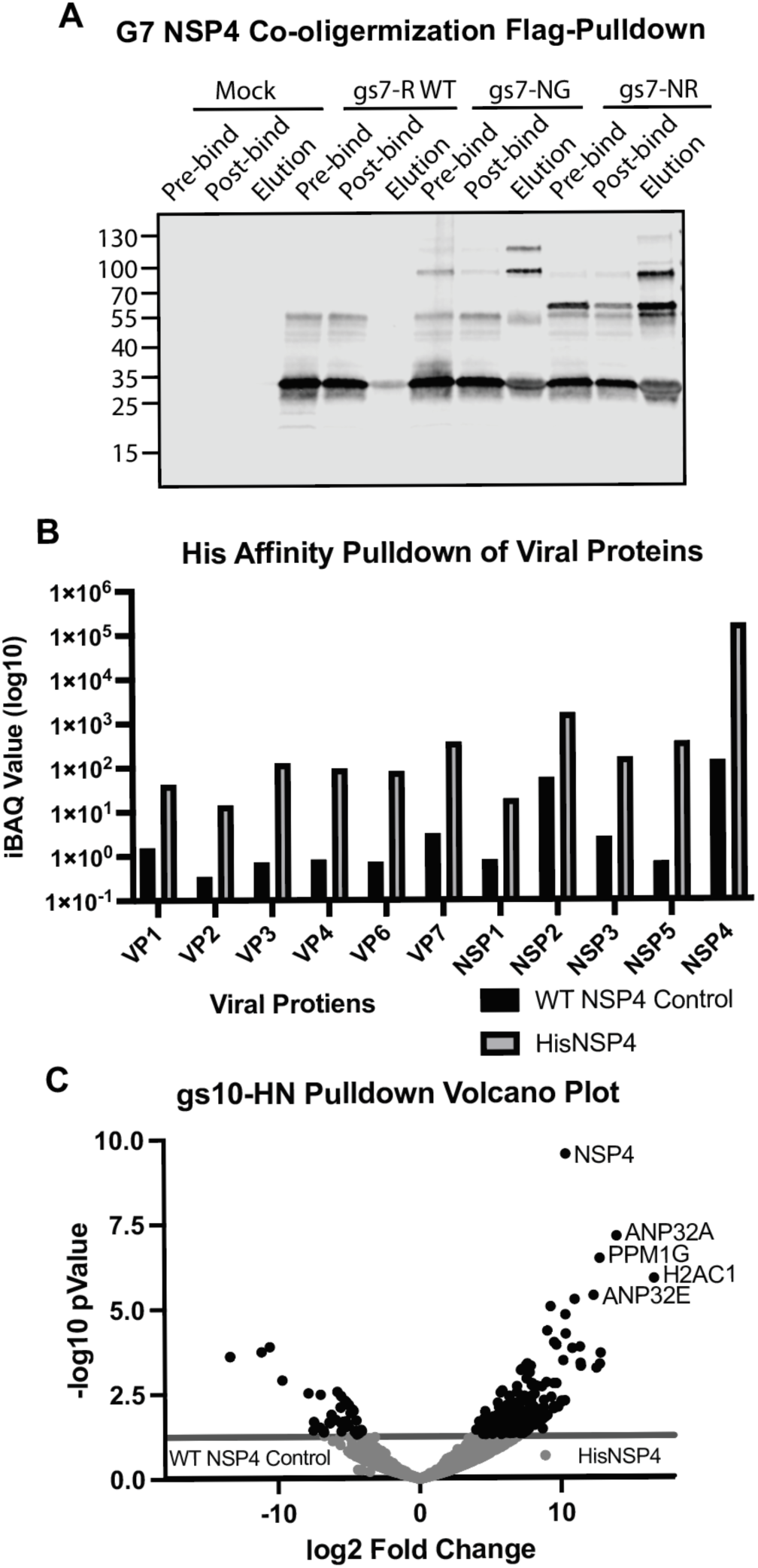
Co-oligomerization and interacting partners of tagged NSP4 during rotavirus infection. (A) FLAG affinity pulldown from cells infected with the indicated gs7 rRV expressing WT NSP4 (gs7-R), gs7-NG, or gs7-NR. Immunoblot analysis with anti-NSP4 aa114-135 antibody and secondary conjugated to IRDye680RD of pre-binding, post-binding, flow-through, wash, and elution fractions demonstrates co-purification of WT NSP4 with FLAG-tagged NSP4 variants, consistent with NSP4 co-oligomerization, N=2. (B) Quantitative mass spectrometry analysis of His affinity-purified proteins from cells infected with the gs10-HN. Relative iBAQ values indicate selective enrichment other viral proteins compared with NSP4. (C) Volcano plot of proteins enriched following His affinity purification from gs10-HN-infected cells relative to the wild-type NSP4 control. NSP4 was the most significantly enriched viral protein.

To characterize proteins associated with NSP4-containing complexes, we purified lysates from rRVs expressing an N-terminal HisTEV-tagged NSP4 from gs10 (gs10-HN) by nickel affinity chromatography and analyzed them by quantitative LC-MS/MS. Furthermore, we assessed additional protein enrichment relative to WT SA11-infected controls. As expected, NSP4 was highly enriched in the His-tag pulldown compared with WT control samples, validating the specificity of the affinity purification (Fig. 6A). In addition to NSP4, several viral proteins co-purified with His-tagged NSP4, including the structural proteins VP7, VP6, VP4, and VP3, as well as the nonstructural proteins NSP2, NSP3, and NSP5 (Fig. 6B). Among these, NSP2 represented the most abundant co-purifying viral protein after NSP4, suggesting a robust association between NSP4-containing complexes and viroplasm-associated proteins.

To identify host proteins interacting with NSP4, we compared proteins enriched in the HisNSP4 pulldown with WT control samples by quantitative proteomic analysis. Volcano plot analysis demonstrated strong enrichment of NSP4, confirming successful purification of the bait protein (Fig. 6C). We also identified several significantly enriched host proteins, including ANP32A, ANP32E, PPM1G, and H2AC1, suggesting that these proteins associate with NSP4-containing complexes during infection. The enrichment of these host factors suggests potential roles in NSP4-mediated replication, protein trafficking, or regulation of viral replication complexes and provides a foundation for future mechanistic studies (13, 21, 32)

Collectively, these data demonstrate that recombinant NSP4 expressed from an engineered genome segment remains capable of co-oligomerizing with WT NSP4 during infection and establish that affinity purification of tagged NSP4 can recover both viral and host proteins associated with NSP4-containing complexes. These findings validate rRVs as useful tools for defining the NSP4 interactome during rotavirus infection.

## Discussion

Despite the availability of RV vaccines, disease burden remains high in low-resource settings, highlighting the need for an improved understanding of the biology and pathogenesis of RV. NSP4, a viral protein with roles in both viral assembly and virulence, is an attractive target to study for RV biology and as a potential therapeutic target. However, studying NSP4 through mutagenesis or modification within infectious virus has remained challenging following the advent of the RV reverse genetics system. To address this, we employed three strategies to modify NSP4 through (1) expression from an alternative gene segment, (2) N-terminal tagging of gs10 by shifting the ORF downstream of the packaging sequence, and (3) retention of the gs10 5’ UTR and first 20 amino acids of the ORF fused to a reporter followed by a bicistronic arrangement to allow full encoding of the NSP4 ORF.

NSP4 functions simultaneously as a viral ion channel and as a structural determinant of virion assembly but remains difficult to modify in the context of a bone fide infection due to gs10 having essential packaging signals at its 5′ end and an indispensable NSP4-DLP-binding domain at its C-terminus. Prior studies have therefore relied predominantly on overexpression systems that lack the spatial and temporal context of active viral replication or relied on the kinetics of other modifiable gene segments. In this study, we used a plasmid-based reverse genetics platform to generate six rRVs expressing tagged or reporter-fused NSP4 from either from an alternative genome segment (gs7) or from an engineered gs10 that preserved critical 5′ and 3′ regulatory elements. These viruses constitute a validated toolkit for interrogating NSP4 structure, assembly, and signaling function during infection.

Genomic structure played a factor in viral fitness, which outweighed the specific insertion applied. All gs7 constructs exhibited delayed replication and for gs7-NR and gs7-NG endpoint yields were approximately 1-log lower than WT SA11. This is consistent with the added burden of bicistronic expression on NSP3 which enhances viral mRNA activity and shuts off cellular protein synthesis. Advances in reverse genetics, including improved codon optimization and sequence engineering, may help mitigate this burden by reducing unfavorable sequence features and improving the genetic compatibility of larger reporter insertions (33, 34). By comparison, all gs10 constructs replicated with similar kinetics to WT SA11 despite more extensive modification of gs10s 5’ architecture. Previous research detected gs10 duplication in circulating RV isolates indicating that increased gene length in gs10 may naturally have greater tolerance (35). The genetic instability of gs7-NG, which exhibited recombination within GCaMP8s after two passages, indicates that larger reporter insertions are poorly tolerated despite short-term replication competence. This barrier for large constructs may exist for other rRV gene segments when utilizing this insertion strategy, necessitating plaque purification and periodic sequence validation.

Co-precipitation of WT NSP4 with FLAG-tagged NSP4 in gs7-NR and gs7-NG pulldowns indicates that rRV gs7 NSP4s assemble into mixed oligomers with endogenous protein rather than forming separate, segregated pools. This supports the physiological relevance of the gs7 based tools but also complicates interpretation of associated phenotypes, since calcium signaling or trafficking readouts in these viruses reflect the combined behavior of tagged and untagged NSP4 populations. However, these methods are unable to resolve stoichiometry of WT-to-tagged species within the oligomers, and whether tagged homomers would be fully functional.

The gs10-N20RN reporter virus enabled live monitoring of NSP4 expression without perturbing downstream signaling. Onset of mRuby fluorescence either preceded or coincided the appearance of the first calcium transients, confirming that reporter expression is a reliable temporal proxy for NSP4 synthesis and enabling single-cell, longitudinal analysis of expression-signaling relationships not readily achievable with fixed endpoint assays.

The inability of gs10-HN to trigger ICWs was a surprising result and as the protein expression kinetics were not decreased, the underlying mechanism for this defect remains unresolved. One possibility is that placement of the HisTEV tag at the N terminus of NSP4, while compatible with viral replication and glycosylation, interferes with one or more specific functional properties of NSP4 required for calcium dysregulation. The N-terminal luminal domain of NSP4 contains the signal peptide and residues involved in ER membrane insertion, oligomerization, and interactions with host factors that regulate calcium homeostasis (10). Although gs10-HN remained Endo H sensitive and formed oligomers, the added HisTEV sequence may alter the oligomeric packing or conformational flexibility of this region, thereby reducing viroporin activity and impairing the ability of NSP4 to stimulate extracellular purinergic release. However, the fact that gs10-N20-HN, which expressed a HisTEV tagged NSP4, triggered ICWs similar to viruses expressing native WT NSP4 indicates that on its own a HisTEV tag does not impair NSP4 viroporin function. The main difference between these viruses is the presence of the mixed population of untagged and HisTEV tagged NSP4 in gs10-HN infections, which was identified in our western blot analyses (Fig 4D, H). While this presents a confounding factor, it is possible that these heterooligomers are responsible for restricting NSP4’s viroporin function. Nevertheless, how the HisTEV tag in a mixed heteromer would restrict the ability of NSP4 to trigger ICWs remains unclear, as the molecular mechanism by which NSP4 viroporin activity triggers purine release to generated ICWs remains to be defined. Additionally, without a high-resolution protein structure of NSP4, the possibility that the HisTEV tag alters NSP4’s conformational flexibility remains purely speculative. Nevertheless, NSP4-mediatd increases in calcium signaling within infected cells is known to be critical for rotavirus replication, so the gs10-HN mixture of native and HisTEVNSP4 retains sufficient viroporin activity to support replication competence but is diminished enough to impair the triggering of ICWs (10, 11, 13, 32, 36, 37). Further experiments measuring ER calcium release, viroporin channel activity, and ADP secretion would be required to distinguish between these possibilities.

Affinity purification of His-tagged NSP4 recovered other viral proteins including VP7, VP6, VP4, VP3, NSP2, NSP3, and NSP5, with NSP2 the most abundant co-purifying viral protein after NSP4 itself. This is consistent with a functional association between NSP4 and viroplasm-associated replication intermediates and harmonizes with the role of NSP4 in coordinating DLP maturation. The joint pulldown of NSP2 and NSP5 indicate that His tag pulldown may precipitate either fully mature or even maturing viroplasms. Among host proteins enriched in the pulldown, ANP32A, ANP32E, PPM1G, and H2AC1 emerge as candidates for future investigation, along with dozens of other significantly enriched host proteins. ANP32 family proteins participate in nuclear export and chromatin regulation in other viral systems, and PPM1G, a serine/threonine phosphatase, could plausibly intersect with NSP4-associated signaling (38–41). These associations remain correlative, and validation by orthogonal methods will be required before mechanistic roles can be assigned. Nevertheless, these preliminary studies establish the utility of this engineered gs10 with both a fluorescent reporter and an affinity tagged NSP4 to examine functional protein-protein interactions during the course of infection. Further, this general engineering strategy should be applicable to different NSP4s, enabling RV strain or NSP4 sequence-based comparisons of protein interaction sites.

This work establishes a reverse genetics strategy that overcomes the packaging and assembly constraints that have historically limited NSP4 mutagenesis, yielding a panel of tools for visualizing NSP4 expression and calcium signaling during infection. The identification of an assembly-competent, signaling-null rRV (gs10-HN) provides a foundation for future studies dissecting the molecular basis of viroporin-associated calcium signaling. The design principle of preserving essential UTR elements while modifying downstream ORF architecture may generalize to other RV gs constrained by similar packaging or assembly requirements.

## Materials and Methods

### Cell lines

MA104 (American Type Culture Collection, CRL-2378.1) cells were cultured in high-glucose Dulbecco’s modified Eagle’s medium (DMEM) with 10% fetal bovine serum and 1x antibiotic/antimycotic (Invitrogen) at 37°C with 5% CO_2_. BHK-T7 (CVCL_RW96) cells were cultured in high-glucose Glasgow minimum essential medium (GMEM) with 5% fetal bovine serum and 1x antibiotic/antimycotic (Invitrogen) at 37°C with 5% CO_2_.

### Reverse genetics

The rRV SA11 strains were generated using established protocols for reverse genetics (24, 42). RVA (simian SA11 strain) plasmids pT7/VP1SA11, pT7/VP2SA11, pT7/VP3SA11, pT7/VP4SA11, pT7/VP6SA11, pT7/VP7SA11, pT7/NSP1SA11, pT7/NSP2SA11, pT7/NSP3SA11, pT7/NSP4SA11, and pT7/NSP5SA11 were provided by Takeshi Kobayashi through the Addgene plasmid repository (24); additionally, pT7/NSP3-P2A-fmRuby was provided by John Patton (25). Recombinant genes synthesized commercially and cloned into the background by epoch life sciences, Missouri City TX.

### Sequencing/RT-PCR

RNA was isolated from viral stocks using a viral RNA extraction kit (Zymo Research) according to manufacturer instructions. cDNA synthesis for gs7 and gs10 was done following manufacturer protocols (PrimeScript™ One Step RT-PCR Kit, Takara Bio). cDNA products were amplified (KOD Hot Start DNA Polymerase, Sigma-Aldrich) and product size was verified by gel electrophoresis. PCR amplification products or gel slices were purified (Wizard® SV Gel and PCR Clean-Up System, Promega). Product yield and quality were assessed on a spectrophotometer (NanoDrop, Thermo Scientific) and sequenced by Sanger sequencing (Azenta). Sequencing contigs were assembled and aligned with SnapGene ver 8.1.1. Primers utilized are listed in Table 1.

### Rotavirus RNA Extraction and Electropherotyping

Viral RNA was extracted from rotavirus-containing samples using the Quick-RNA Viral Kit (Zymo Research, Cat. No. 11-355B) according to the manufacturer’s instructions.

Rotavirus genomic RNA segments were separated by agarose gel electrophoresis. RNA samples were mixed with RNA loading buffer and loaded onto a 1% agarose gel prepared with 1x TAE containing ethidium bromide. Electrophoresis was performed in 1x TAE at 120V at room temperature. Following electrophoresis, gels were visualized using a UV transilluminator. Rotavirus genomic electropherotypes were determined based on the characteristic migration pattern of the 11 gene segments. Segment migration patterns were compared between recombinant viruses and the parental SA11 strain to confirm the expected genome profile and assess the presence of altered RNA segments.

### One step Growth Curve

Confluent MA104 cell monolayers were infected with the indicated strains of RV for 1 hour at an MOI of 5. The inoculum was removed, and monolayers were washed with once with 1X PBS before replacement with serum-free high-glucose DMEM. At 2, 4, 8, 18, and 24 hpi, monolayers underwent three freeze-thaw cycles, were transferred to 1.5-ml microcentrifuge tubes, centrifuged to remove cell debris, and incubated for 45 min at 37°C with Worthington’s trypsin (10 μg/ml). RV yield was assessed by FFA as described below.

### Imaging and antibodies

Immunofluorescence staining and imaging of monolayers were conducted using established protocols with the indicated primary and fluorescently conjugated secondary antibodies (Table 2) (13).

**Table 2.**
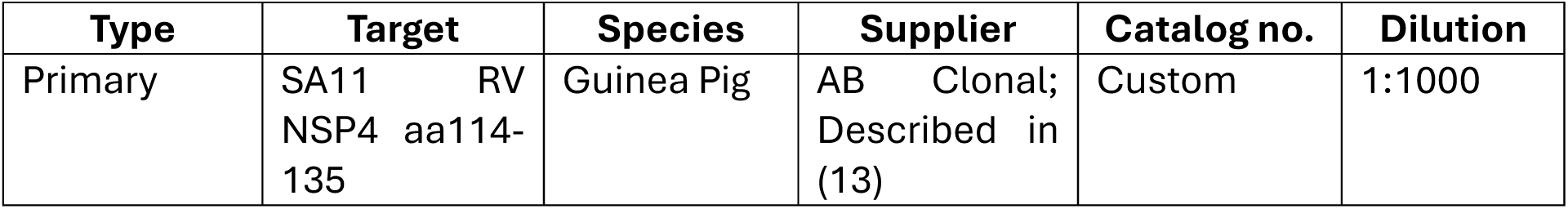
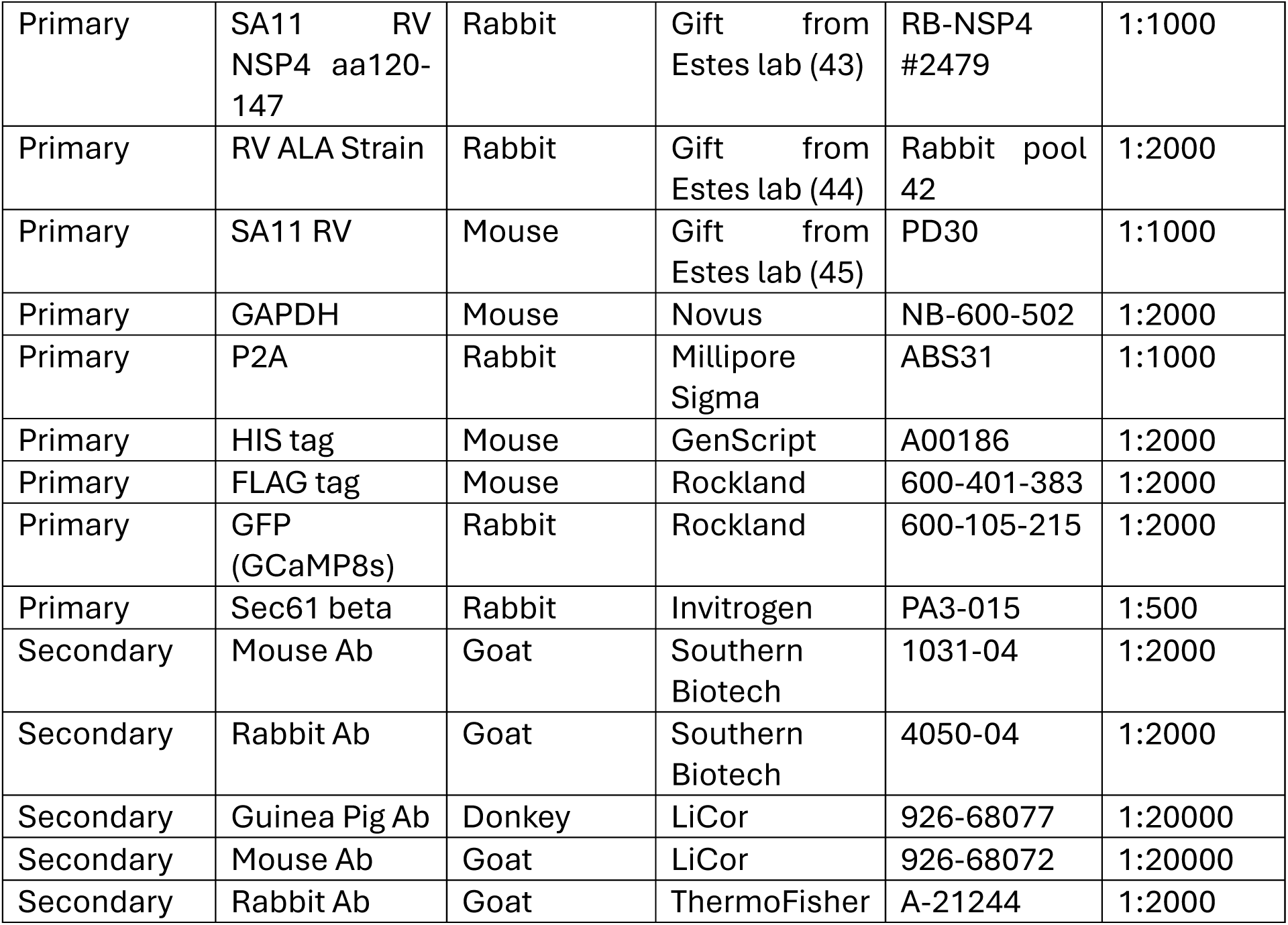
Antibodies.

### Solid Overlay Plaque Picking

RV infected lysates (cells and supernatant) underwent three freeze-thaw cycles and activation with Worthington’s trypsin (10 μg/ml) for 30 min at 37°C before 10-fold serial dilution. MA104 cells were grown to confluency in six-well plates and switched to serum-free medium for 24 hours before infection. Wells were infected with 200 μl of each serial dilution in duplicate for 1 hour at 37°C with gentle shaking every 15 min to ensure even distribution. Monolayers were rinsed once with 1x PBS, and medium was replaced with a solid overlay consisting of 1.2% Seakem Agarose in serum-free DMEM supplemented with O-(diethylaminoethyl) dextran and Worthington’s trypsin (1 μg/ml). After 72 hours, cells were imaged with using a 10× Plan Apo objective with agarose plugs taken of fluorescent positive plaques.

### Nickel Affinity Purification of His-Tagged Proteins from Rotavirus-Infected MA104 Cells

Confluent mock- or rotavirus-infected MA104 monolayers in T150 flasks were lysed in 2.5 mL M-PER™ (Thermo Scientific) per flask, pooled, and clarified by centrifugation (4000 rpm, 10 min, 4°C). Supernatants were collected without disturbing pellets. Ni-NTA agarose beads (Qiagen; 250 µL bead volume) were equilibrated by two washes with PBS + 10 mM imidazole and one wash with M-PER + 10 mM imidazole (centrifugation at ∼13,000 rpm, 1 min, 4°C per wash). Beads were incubated with 1 mL lysate + 10 mM imidazole at 4°C for ≥2 h with end-over-end mixing, then pelleted (2500 × g, 5 min, 4°C) and the flow-through was saved. Beads were washed twice with 1 mL M-PER + 10 mM imidazole (10 min, 4°C, mixing), pelleted, and washes collected. Elution was performed twice with 200 µL CHAPS elution buffer (0.5% CHAPS, 500 mM imidazole in M-PER), incubated 10 min at 4°C with mixing, pelleted, and eluates saved. Beads were then resuspended in 1 mL elution buffer, and 50 µL was mixed with 10 µL 5x SDS sample buffer to generate the beads post-elution (BPE) fraction.

### Mass Spectrometry

The BCM Mass Spectrometry Proteomic Core ran provided pulldown samples on the 1D SDS-PAGE, cut bands, and in-gel digested the proteins and sequenced 2 pooled peptides fractions per sample on a Orbitrap mass spectrometer. The data analyzed against *Chlorocebus sabaeus* (NCBI RefSeq database 60711) with gpGrouper (46). The core performed evaluation of quality control (QC) metrics, principal component analysis (PCA), clustering (hierarchical, k-means), differential expression (volcano plots and statistics), gene-set enrichment analysis (GSEA), and annotation of relevant protein categories of interest.

### Anti-Flag Tag Magnetic Pulldown

Confluent mock- or rotavirus-infected MA104 monolayers in T75 flasks were lysed in 1 mL M-PER™ (Thermo Scientific) per flask and clarified by centrifugation (4000 rpm, 10 min, 4°C). Anti-Flag agarose beads (Qiagen; 50 µL bead slurry volume) were incubated with 1 mL lysate at 4°C for ≥48 hours with end-over-end mixing, then magnetically pelleted (5 min, RT) and the flow-through was saved. Beads were washed 4 times with 1 mL 1x PBS (10 min, mixing), and pelleted. Elution was performed with 60 µL 1x SDS sample buffer in M-PER, incubated 10 min and saved.

### Western blot

Lysate generation, gel electrophoresis and membrane transfer was conducted as previously described (13). Alkaline phosphatase (AP) western blots were conducted as previously described (Table 2) (47). Due to the successful rescue of superior His tagged NSP4 RVs we only displayed the gs7-NH 10 hpi time point to demonstrate viral protein expression. Fluorescent western blots were conducted as previously described (Table 2) (13).

### Calcium imaging

MA104 cells stably expressing the cytosolic calcium indicator GCaMP6s were generated using lentivirus transduction as previously described (48). Monolayers were seeded in eight-well, imaging-bottom chamber slides (Ibidi) and allowed to grow until confluent. Monolayers were mock or RV infected at a MOI of 0.05 for 1 hour. The monolayers were then imaged as previously described by (13, 49).

### Image analysis

For the ICW analysis of image series from long-term calcium imaging experiments, we used the open-source platform FIJI as previously described (13, 49, 50).

### Semi-solid Overlay Plaque assay

Plaque assays were performed as previously described (12, 13).

### Fluorescence focus assay

Fluorescent focus assays were prepared, infected, stained, and imaged as previously described (13). Quantification of the number of infected cells per well was conducted via manual eye count.

## Acknowledgements

This work was supported by National Institutes of Health (NIH) grants NIAID R01AI158683 and NIDDK R01DK115507 to J.M.H., S.E.C. and M.K.E, NIH grant F30AI169983 to J.L.P., NIH grant T32GM136554 to Ignatia Van Den Veyver and Melissa Suter partially supporting E.M.H. BCM Mass Spectrometry Proteomics Core (RRID:SCR_027015) is supported by the Dan L. Duncan Comprehensive Cancer Center Award (P30 CA125123), CPRIT Core Facility Awards (RP210227), Intellectual Developmental Disabilities Research Center Award (P50 HD103555), and NIH High End Instrument Award (S10 OD026804, Orbitrap Exploris 480).

---

**Supplemental Figure 1.**
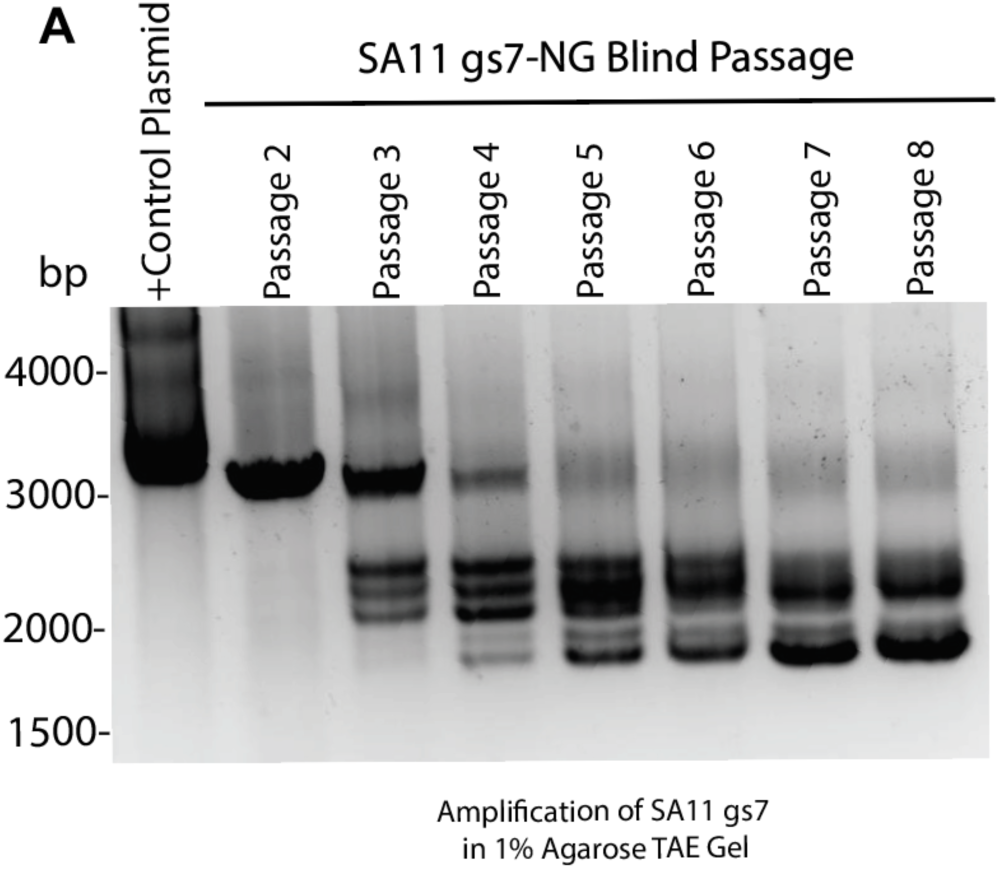
Genetic stability of reporter-tagged rotavirus genomes during blind passage. (A) RT-PCR amplification of the SA11 gs7-NG genome segment following serial blind passage. The gs7 amplicon pattern changed beginning at passage 3, with the appearance of multiple lower-molecular-weight products that persisted through passage 8, consistent with instability of the gs7-NG reporter-containing genome segment. (B) RT-PCR amplification of the SA11 gs10-N20RN genome segment following blind passage. The predominant amplicon remained at the expected size across passages 2–10, although additional lower-molecular-weight products were observed at passage 8. PCR products were resolved on 1% agarose gels alongside molecular weight markers.

## Notes

### Competing Interest Statement

The authors have declared no competing interest.

